# Insects and recent climate change

**DOI:** 10.1101/2020.03.09.984328

**Authors:** Christopher A. Halsch, Arthur M. Shapiro, James A. Fordyce, Chris C. Nice, James H. Thorne, David P. Waetjen, Matthew L. Forister

## Abstract

Insects have diversified through 400 million years of Earth’s changeable climate, yet recent and ongoing shifts in patterns of temperature and precipitation pose novel challenges as they combine with decades of other anthropogenic stressors including the conversion and degradation of land. Here we consider how insects are responding to recent climate change, while summarizing the literature on long-term monitoring of insect populations in the context of climatic fluctuations. Results to date suggest that climate change impacts on insects have the potential to be considerable, even when compared to changes in land use. The importance of climate is illustrated with a case study from the butterflies of Northern California, where we find that population declines have been severe in high-elevation areas removed from the most immediate effects of habitat loss. These results shed light on the complexity of montane-adapted insects responding to changing abiotic conditions and raise questions about the utility of temperate mountains as refugia during the Anthropocene. We consider methodological issues that would improve syntheses of results across long-term insect datasets and highlight directions for future empirical work.

**Significance statement:** Anthropogenic climate change poses multiple threats to society and biodiversity, and challenges our understanding of the resilience of the natural world. We discuss recent ideas and evidence on this issue and conclude that the impacts of climate change on insects in particular have the potential to be more severe than might have been expected a decade ago. Finally, we suggest practical measures that include the protection of diverse portfolios of species, not just those inhabiting what are currently the most pristine areas.

---

From invasive species to habitat loss, pesticides and pollution, the stressors of the Anthropocene are many and multi-faceted, but none are as geographically pervasive or as likely to interact with all other factors as climate change (1, 2). For these reasons, understanding the effects of anthropogenic climate change on natural systems could be considered the defining challenge for the ecological sciences in the 21^st^ century (3). It is of particular interest to ask how insects will respond to recent and ongoing climate change, because they are the most diverse lineage of multicellular organisms on the planet, and of fundamental importance to the functioning of terrestrial ecosystems. The issue also has new urgency in light of recent and ongoing reports of insect declines from around the globe (4). Insects and climate change have been discussed elsewhere (5–8), and our goal here is not to cover all aspects of the problem. Instead, we focus on recent discoveries and questions inspired by long-term records of insect populations, discussing other fields (e.g., physiological ecology) where they inform our understanding of population trajectories under climate change scenarios.

In the sections below, we compare climate change to other stressors and examine multifaceted impacts in terms of climate means, limits and extremes. Then we discuss the geography of climate change with particular focus on the responses of montane insects, with a case study from the butterflies of Northern California that illustrates the value of long-term observations that span a major gradient of land use intensity. Two areas that we do not cover in detail are the theoretical foundations of climate change research (9), and community-level consequences including altered trophic interactions associated with shifting phenologies (10). As a qualitative survey of the state of the field, we have gathered insect monitoring studies (based on a review of more than 2,000 papers) that encompass at least 10 years of continuous sampling and analyses of climatic data (Table 1). We excluded studies of pest species and studies from agriculture, as well as projects encompassing fewer than 10 species (as a way to focus on assemblages of species rather than model systems). In general, we emphasize monitoring studies as unparalleled opportunities for understanding the influence of climatic fluctuations on animal populations because of the ability to decompose complex temporal trends into effects driven by different factors (11, 12).

**Table 1.** Monitoring studies of at least 10 insect species and at least 10 years from land use gradients or protected areas that have been used to examine weather in relation to insect populations.

| Location | Source | Years | Species | Taxa | Method |
| --- | --- | --- | --- | --- | --- |
| Australia | Gibb et al. (58) | 21 | 106 | Ants | Pitfall traps |
| California, USA | Shapiro Transect (27, 41–43) | 47 | 163 | Butterflies | Modified Pollard walk |
| Colorado, USA | Iler et al. (59) | 20 | 20 | Hoverflies | Malaise traps |
| Costa Rica | Tritrophic Interaction Monitoring in the Americas (31) | 22 | 1724 | Lepidoptera, Parasitoids | Collect and rear |
| Ecuador | Grøtan et al. (53) | 10 | 137 | Butterflies | Fruit traps |
| Europe | Jourdan et al. (60) | 32 | -- | Benthic invertebrates | Surface water survey |
| Finland | Finnish Moth Monitoring Scheme (61) | 13 | 183 | Moths | Light traps |
| Finland | Hunter et al. (23) | 32 | 80 | Moths | Light traps |
| Germany | Voight et al. (62) | 20 | 1041 | Arthropods | Pitfall trap, Sweep net |
| Germany | Krefeld Entomological Society (63) | 27 | -- | Flying insects | Malaise traps |
| Germany | Baranov et al. (22) | 42 | 125 | Mayflies, stoneflies and caddisflies | Emergence trap |
| Great Britain | National Moth Recording Scheme (20) | 40 | 673 | Moths | Traps |
| Greenland | Koltz et al. (29) | 18 | -- | Arthropods | Pitfall traps |
| Netherlands | Dutch Monitoring Scheme (33, 64) | 18 | 40 | Butterflies | Pollard walk |
| Netherlands | Hallmann et al. (65) | 28 | -- | Beetles, Moths, Caddisflies | Pitfall traps |
| Russia | Chronicles of Nature (66) | 40 | 19 | Arthropods | Traps |
| Spain | Stewart et al. (67) | 10 | 10 | Butterflies | Pollard walk |
| United Kingdom | UK Butterfly Monitoring Scheme (32, 34, 54, 68–73) | 45 | 55 | Butterflies | Pollard walk |
| United Kingdom | Rothamsted Insect Survey (28, 32, 54, 74) | 50 | 345 | Aphids, Moths | Light trap |
| United Kingdom | Hoverfly Recording Scheme (75) | 54 | 215 | Hoverflies | Citizen observations |

## On the relative importance of climate change and other stressors

Although Anthropocene stressors must ultimately be understood as an interacting suite of factors (13), it is useful to start by asking: how will the consequences of climate change compare to other stressors? Over the last three centuries, the global percentage of ice-free land in a natural state (not intensively modified by human activity) has shrunk from 95% to less than 50% (14), with consequences that include the extirpation and extinction of plants and animals (15). Although habitat loss (including degradation through pollution and numerous other processes) continues, it is possible that we are living through a period of transition where the importance of changing climatic conditions could begin to rival the importance of habitat loss (16, 17). That transition and the rise of climate change as a more severe threat could happen through a number of related processes. A very partial list would include the inability to disperse quickly enough to track changing conditions (18), or the phenomenon of dispersal out of previously suitable habitat into regions where habitat is suboptimal or cannot be found at all (19). There is also the possibility that abiotic stress associated with changing climatic conditions could simply be the last straw, reducing vital rates below replacement levels as a result of physiological stress in populations already pushed to the edge by other stressors (5).

An empirical understanding of the effects of climate change in comparison to other stressors depends in large part on long-term observations from protected areas or from gradients of land use that will let us directly compare the effects of different factors. In Great Britain, both land use and climate change have been important for explaining the decline of 260 species of macro-moths and an increase of 160 species (out of a total of 673 species) (20). The signal of habitat loss is seen in widespread species, which have declined in regions with increased intensity of human land use. At the same time, the role of climate can be seen in the decrease of more northern, cold-adapted species and the simultaneous increase of more southern, warm-adapted species (20). Less multifaceted signals of global change can be found in smaller areas sheltered from direct effects of habitat loss. Beetle activity in a protected forest in New Hampshire has, for example, decreased by 83% in a re-sampling project spanning 45 years, apparently as a function of warmer temperatures and reduced snow pack that insulates the diverse ground-dwelling beetle fauna during the coldest months (21). In a headwater stream in a German nature preserve that has been isolated from other anthropogenic stressors (besides climate change), abundance of macroinvertebrates has declined by 82% over 42 years and mean emergence is more than two weeks earlier, while species richness has increased (22). It is important to note that a signal of climate has not been found in all long-term studies of insects, even those from protected areas. In a sub-arctic forest in Finland, moth populations are primarily stable or increasing and these trends do not appear to be strongly related to warming temperatures (23). It can also be noted that the literature on long-term responses of insect populations to climate is neither deep nor geographically broad, which is an important conclusion from Table 1, where it can be seen that most studies come from Northern Europe and Lepidoptera are disproportionately represented, as others have noted (4).

Beyond the direct effects of climate change, we can ask: how will changing climatic conditions interact with habitat loss, invasive species, pesticide toxicity (24), and other factors? This is an area that is ripe for experimental work (11), but the number of potentially interacting factors that could be tackled in an experiment is daunting, which is why experiments will profitably be inspired and focused by observational results. The challenge for researchers with long-term data is to develop statistical models that encompass interactions rather than focusing only on main effects that might be easier to interpret. A notable example of modeling interactions in the context of global change comes from a recent study of British insects, where researchers found that the most successful model for poleward range shifts included habitat availability, exposure to climate change, and the interaction between the two (25).

Our reading of the literature comparing climate to other drivers of change suggests a few methodological issues that could be better aligned across future studies. Results from analyses of weather and insect populations should be reported as standardized beta coefficients, and summarized at both seasonal and annual scales; finer scales may be appropriate for certain questions or datasets, but those two broader scales would facilitate comparisons among studies. Whenever possible, year or time as a variable should be included in models with weather explaining insect population or community data. Conditioning on year strengthens the inference of causation, especially when variables (insects and climate) are known *a priori* to be characterized by directional change. When year cannot be included in models because of high collinearity with weather or other variables, first order differencing or other methods of trend decomposition should be considered (12).

## On changing maximums, minimums, means and variance

Climate change is of course not one phenomenon, and axes of change include shifts in limits (maximums and minimums), average conditions and variance, which can all be measured at different temporal and spatial scales. The multifaceted nature of climate change is illustrated by the fact that nighttime temperatures are warming faster than daytime conditions (26), and the consequences for insects are poorly-understood but potentially serious, including reduced time for recovery from daytime heat stress, and indirect effects through plants (26). In the mountains of California, rising average daily minimum temperatures had some of the most dramatic negative effects on insects, especially in combination with drier years (27). Rising minimum temperatures in particular seasons might impact insects through effects on critical overwintering stages. In the UK, the annual population dynamics of moths are affected by overwintering temperature and precipitation (28). In this case, winter precipitation has a negative association with moth abundance, while winter temperature has a positive association (28). In Greenland, changes in the structure of arthropod communities over 18 years have been influenced by warming summers and falls and fewer freeze-thaw events, with the most negative associations observed for surface detritivores (29). On the other side of the temperature spectrum is maximum temperature, which has been shown to be the variable most associated with local extinctions in a global survey of insects and other taxa (18).

While our understanding of biotic response to warming means and limits improves, the greater challenge of changing variance is now upon us: predictions for many parts of the world include an increased frequency of extreme weather, which can appear as more intense precipitation events separated by more prolonged dry periods (30). We have few studies on this topic from which to draw conclusions; only five studies in Table 1 explicitly investigated extreme weather events (27, 31–34). In the few cases where biotic response to extreme events has been examined, the results are as we might expect: extreme events are extreme population stressors. Large, synchronized population swings of Lepidoptera in the UK are associated with extreme climate years and responses to these years were negative in 5 out of 6 cases (32). On a continental scale, a recent re-survey of 66 bumble bee species across two continents points to temperatures outside of historical ranges as a major driver reducing occupancy across the landscape (35). Salcido et al. (31) report an increase in extreme flooding events as one of the factors contributing to the loss of parasitoids and Lepidoptera in a Costa Rican forest, which includes the disappearance of entire genera of moths (minimum temperatures also had strong negative effects, consistent with other results discussed above).

## On the geography of biotic response to climate change

An important test of our understanding of ectotherm response to abiotic conditions is the extent to which we can understand and predict responses of insects living in different biomes or climatic regions (36). It has been suggested that tropical insects are more sensitive to warming conditions because they are already at their optimal temperatures, and are thus closer to detrimental thermal maxima relative to temperate insects (37); this is, however, an area of ongoing investigation (38). A related issue is the effect of climate change along elevational gradients, and at least a few expectations align to suggest that montane insects could fare better in climate change scenarios as compared to insects in less topographically complex environments (39). First, montane insects have the opportunity for upslope shifts in range, and the tracking of similar environmental conditions in space is potentially the best buffer against changing conditions. Second, montane insects have access to a greater diversity of microhabitats, which could allow for behavioral thermoregulation even without changes in elevational range. Third, relative to lowlands that are degraded in many parts of the world (because of the concentration of agriculture or urban areas), insects on mountains will often find a greater diversity of plant resources, which (at least for herbivorous insects) should provide some buffer against climate-induced changes in the plant community. Are these expectations borne out by long-term records of insect populations? The answer to that question has applied relevance because it affects how we think about land protections, and whether or not mountains can be climate refugia during the upheavals of the Anthropocene (40).

Few insect monitoring programs encompass extensive elevational gradients, but one exception is the Shapiro Transect across Northern California: ten sites and 163 species of butterflies over more than 2500m of elevation, including a severe gradient in land use, from the intensely modified Central Valley to above tree line in the Sierra Nevada (Fig. 1A and B). Observations have been taken every two weeks during the butterfly flight season for between 32 and 48 years, depending on the site; details of data collection have been described elsewhere (41–43).

**Fig. 1.**
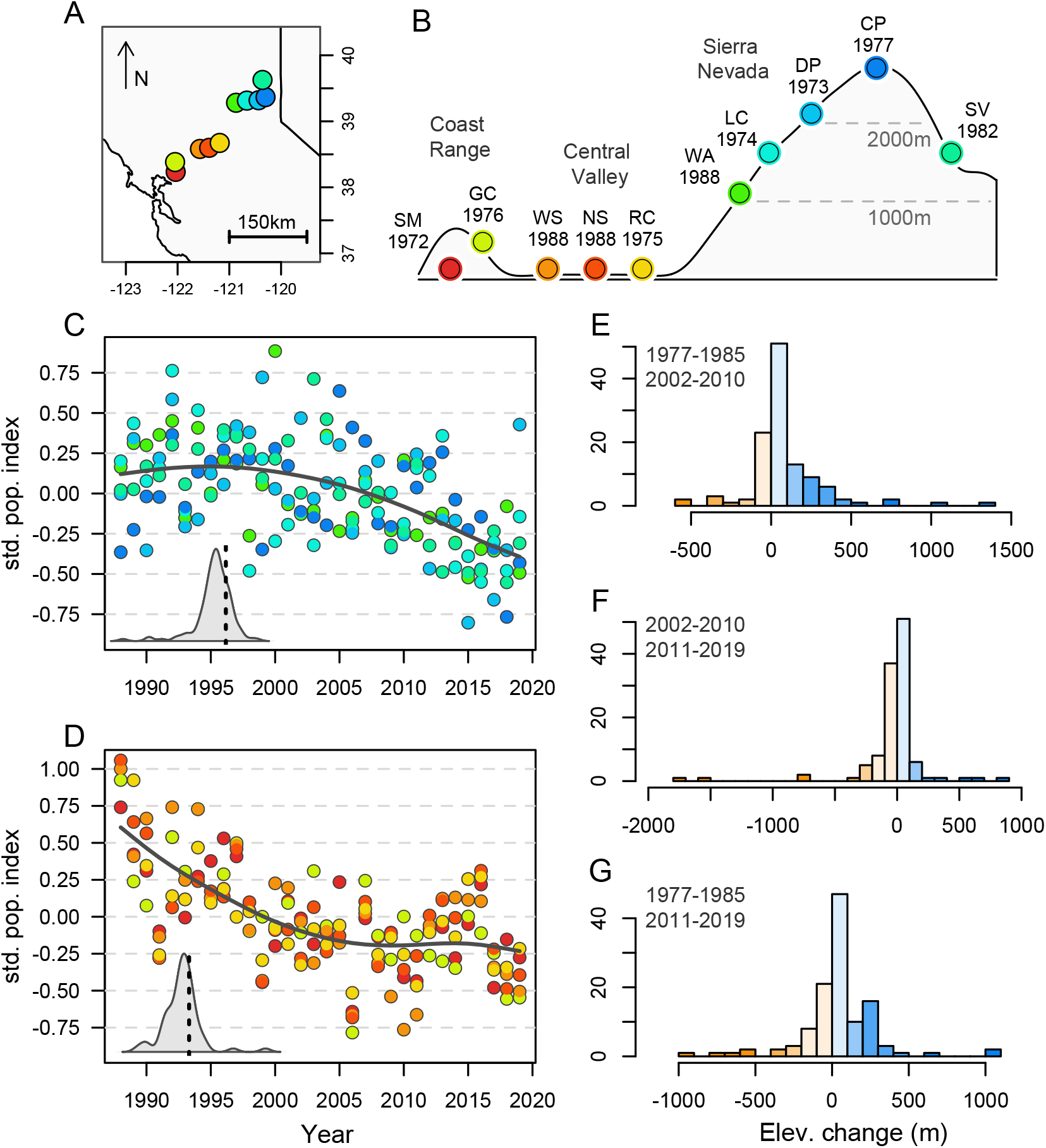
Overview of geography and major trends for Northern California case study. (*A*) Map of Northern California with focal sites, also shown in elevational profile in (*B*), with two-letter site abbreviations and the year when continuous sampling started at each site. (*C* and *D*) Standardized population indices (means across species by site) for mountain sites (*C*) and low elevation sites (*D*), with site colors the same as in (*B*). Inset density plots in (*C* and *D*) show the distribution of year coefficients across species in the two regions (high and low elevations), with vertical dotted lines marking zero, such that observations to the left of the line represent species with negative trends across time. (*E* - *G*) Histograms summarize changes in elevation between different nine-year windows of time; for example, panel (*E*) is the change in mean elevation per species between the earliest years (1977-1985) and years immediately before the mega-drought (2002-2010). Colors in histograms are for visualization with darker orange corresponding to more negative (downward) shifts and darker blue being more positive (upslope) shifts (see *SI Appendix*, Fig. S1 for additional details).

Previous modeling work has highlighted the complexity of population response to weather in this diverse fauna (41), and has documented the array of factors impacting populations along the elevational and land use gradient. At lower elevations, the loss of open spaces, warming summers and pesticide application have been associated with widespread declines (42, 44), while the impact of climate change and an extreme drought have been more apparent at higher elevations (27). Here we revisit the question of climate change impacts in this system (with an additional three years of data), with an emphasis on understanding species-specific traits that predict persistence in the mountains. We also revisit a previously-described upslope shift (43) with an additional 13 years of data to ask if elevational dynamics were impacted by the mega-drought of 2011-2015.

Montane and valley populations have, on average, followed downward trajectories (Fig. 1C and D). Populations at low elevations have been trending downward for a greater span of years, while montane populations appear to have been relatively steady through the 2000s, but were severely impacted following 2011 (the start of the drought years). Roughly speaking, this comparison is between populations affected by all of the major Anthropocene stressors (in the Central Valley) and populations affected primarily by a changing climate (in the mountains). The mountains are not without some land conversion and incursions of invasive plants along roadways, but for the most part our sites are in undisturbed natural areas. Thus, it is noteworthy that the montane declines have reached almost one half of a standard deviation away from the mean (relative to the long-term average), matching roughly the depth of decline in the valley.

The inset density plots in Fig. 1C and D reflect the distribution of demographic trends in the two regions: the bulk of coefficients (associated with years in multiple regression models) are negative (reflecting downward trajectories) in both cases, but not all species are in decline. With respect to the mountains, it is of interest to ask if species with better performance are species that have been observed with greater frequency at the highest elevations, which would be consistent with a bioclimatic (upslope) niche-tracking model. We have updated (in Fig. 1E) an earlier analysis (43) from before the mega-drought years, and confirm that butterflies were on average being observed at slightly higher elevations in later (2002-2010) vs earlier (1977-1985) years; the distribution of those elevation changes in Fig. 1E is positive and upslope (*t* = 3.82, df = 116, *p* < 0.001). A shift in average elevation of occurrence (or change in central tendency of elevational range) is consistent with vegetation dynamics observed in another California mountain range (45). In contrast, when the early vs late comparisons encompass the drought years in a recent (Fig. 1F) or broader span of years (Fig. 1G) it can be seen that the elevational changes are more evenly balanced with both upslope and downslope shifts. This is not unlike the complexity of upslope and downslope responses observed in other taxa in the same mountain range (46, 47).

The severe declines of the drought years in Northern California have in effect cancelled out the earlier upslope signal, which leaves us with the question of whether or not success (or failure) in the mountains in recent years can be predicted based on species-specific traits. We took a constrained ordination approach (redundancy analysis) to understand montane butterfly populations over time in the context of potential predictors that include voltinism (number of generations per year), habitat association, overwintering biology, sensitivity to specific weather variables, and other traits. Focusing on the west slope locations (relevant to our measure of elevational population dynamics in Fig. 1 E-G), we see that the most successful montane species can be characterized as mostly resident (reproducing at our sites), univoltine species with earlier emergence and positive responses to precipitation and average minimum daily temperatures (Fig. 2). The converse is that declining montane species (in the lower half of Fig. 2) have a negative association through time with minimum temperatures, which is consistent with a previous analysis, focused on species richness (27), that hypothesized rising minimum temperatures as a driver of declining montane butterflies. The association with precipitation sensitivity suggests that a successful subset of the montane fauna not only persists with warming nights but is able to take advantage of the highly variable precipitation of the region (27).

**Fig. 2.**
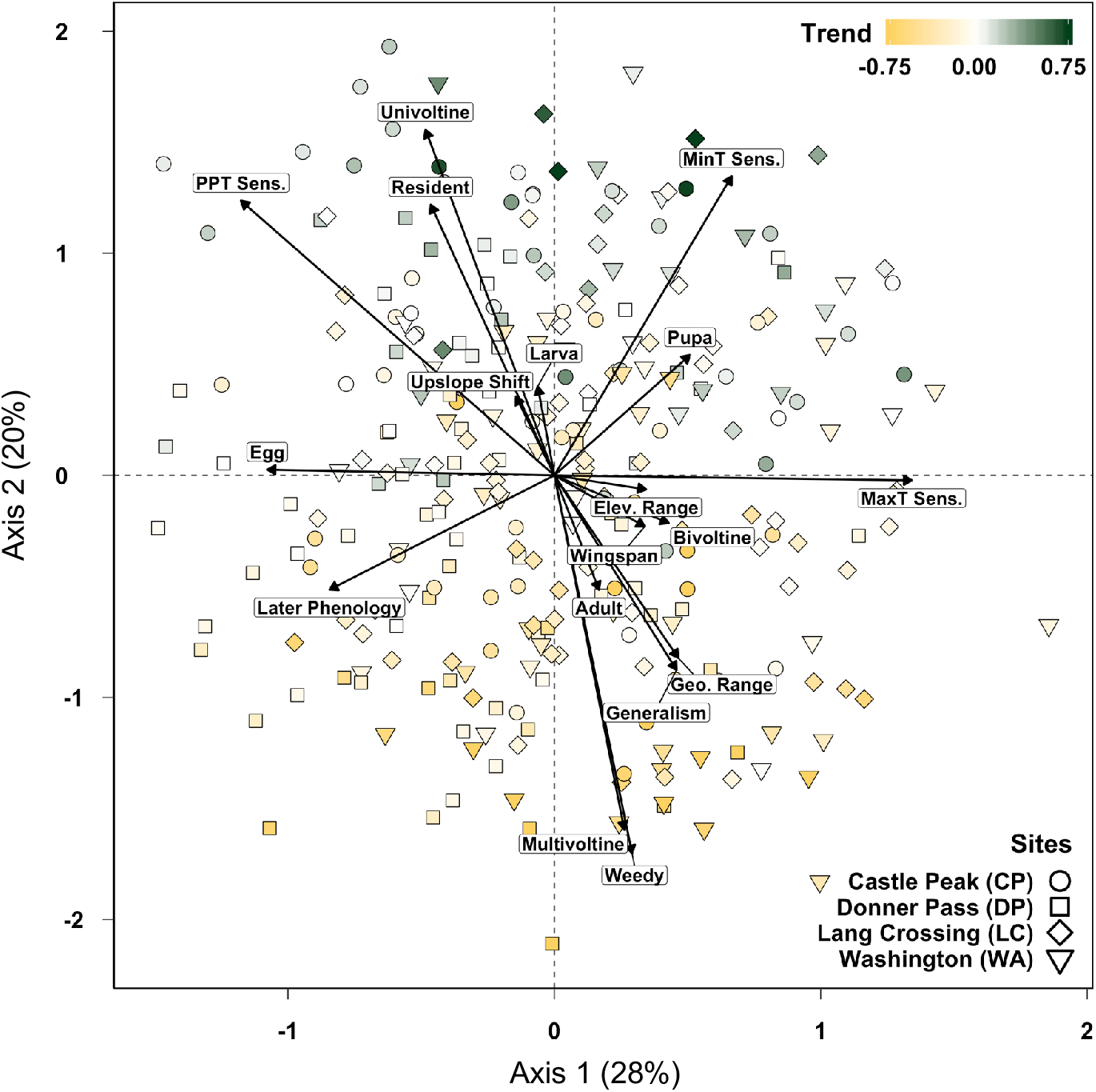
Montane populations through time and population-specific properties that include life history traits and sensitivities to climatic variables, specifically precipitation (PPT Sens.), average daily maximum temperatures (MaxT Sens.) and average daily minimum temperatures (MinT Sens.). For example, populations in the upper portion of the ordination are characterized by positive responses to years that are wetter and have warmer minimum temperatures. Points are colored according to coefficients associated with years (i.e., “trend” or change through time), and those coefficients as well as climate sensitivities were estimated in separate Bayesian models. Each point in the ordination is a population (a species at a site), thus individual species are represented by between 1 and 4 points (depending on their presence at the four mountain sites). Life history traits include overwintering states, geographic range, phenology (average date of first flight), elevational range, elevational shift (as in Fig. 1 E-G), voltinism, body size (wingspan), breadth of habitat association (generalism), and weedy status (see main text and *SI Appendix* for more details). Percent of constrained variation explained is shown in parentheses after each axis label.

Declining populations in the mountains (in the lower half of Fig. 2) tend to be weedy, multivoltine habitat generalists with broad geographic ranges. This result is perhaps superficially surprising given the resilience of generalist species in other contexts (48, 49), but was predicted ten years ago for the montane Northern California fauna (43), and has been seen for multivoltine butterflies in another seasonally hot and dry Mediterranean climate (50). For most species, the warm season at higher elevations is not long enough to support true multivoltinism (51), thus species with many generations per year depend on demographic contributions from lower elevations, where populations have been failing for at least two decades (Fig. 1D). It is interesting to note that having multiple generations per year, however, conveyed the opposite effect at the lowest elevations during an extreme weather event, where we observed that multivoltinism combined with early springs allowed valley populations to reach higher densities during the drought years of 2011-2015, which can be seen in Fig. 1D (27).

These results, which encompass between 100 and 142 butterfly species (depending on the analysis), challenge some of the expected patterns of biotic response to climate change. First, montane microclimatic heterogeneity might not be a strong buffer against climate change. Declines in the mountains are comparable to declines in the Central Valley, which is experiencing other anthropogenic stressors and contains less climatic variation over short distances. These results highlight the power of long-term data to quantify climate sensitivities along with natural history when understanding population trajectories under climate change. These results also bring into focus the complexities faced by organisms when traits (such as voltinism) confer different advantages and disadvantages at locations that are potentially within dispersal distance but separated by elevational, climatic, and habitat differences.

## Conclusions and practical lessons

Reports of insect declines in the scientific and popular press have been greeted with surprise, which could reflect an inherent bias that even scientists have towards assuming that the smallest and most diverse animals on the planet would somehow be more robust than, for example, amphibians or birds. Insects have, after all, seen more than 400 million years of climate change. Can recent and ongoing climate change be that different from others that insects have weathered? In previous periods of change, we know from the paleontological record that individual beetles have relocated across continents (52). As can be seen from Table 1, there are relatively few studies where long-term records of insect populations (with 10 or more continuously-sampled years) have been analyzed in the context of climatic fluctuations. Even more important, only two of those studies are from tropical areas (31, 53), where the majority of insects live, which thus represents a major gap in our understanding of terrestrial biodiversity in the Anthropocene. Nevertheless, considering results from the studies in Table 1 along with spatial or occupancy surveys (e.g., 18), conclusions do emerge. Ongoing climate change will have positive effects on some species and negative effects on others (54, 55), with the balance (of positive and negative effects) determined in some cases by geographic factors such as latitudinal position (20, 37) and in other cases by more complex species-specific traits (6, 7), as in the Northern California case study (Fig. 2). Extreme weather events or prolonged stretches of weather outside of historical conditions will have more consistently negative effects across species (4, 56), although this in an area where additional research is urgently needed.

Moreover, the combination of climatic effects with habitat degradation will certainly have interacting consequences (34, 43), which leads to the conclusion that the current crisis is indeed different than previous periods of Earth history, for the reason that the planet has changed in so many other ways as a result of the increasingly rapid conversion and loss of natural resources associated with the Anthropocene (13). The modernization of agriculture has removed the weedy edges, previously open land has been paved (42), and prolonged droughts have compressed and fragmented tropical cloud forests (56). Nevertheless, we believe that the study of long-term insect records offers some tangible hope and practical lessons. In all but the most severe cases, there are some species that manage to take advantage of anthropogenically-altered conditions (55). Unlike animals with larger home ranges and greater per-individual resource requirements, insects are remarkable in the speed with which they respond to a bit of hedgerow improvement or even a backyard garden. In our own experience, we have been surprised by the resilience of the low elevations of Northern California (27). Some of these places are far from land that you might spot as a target for protection: rights of way, train tracks, levees, or drainage ditches. Yet it was the butterflies in those places that proved to be the most robust during the mega-drought. In the mountains, we have reported success conferred by combinations of traits that could only have been partly predicted by previous work. Of course, the butterflies at low and high elevations in California still continue a downward trajectory of which climate plays no small part, but if other stressors could be alleviated it might be the case that insects even in close proximity to human development will continue to do what insects do best: survive.

## Methods

The literature search was performed on ISI Web of Science in February 2020 using the search terms TS=(insect* OR lepidoptera* OR hymenoptera* OR diptera* OR hemiptera* OR coleoptera*) AND TS=(climate OR weather) AND TS=(“long term” OR “long-term” OR monitor*), which identified 2,264 studies. To be included in Table 1, we considered studies that included at least 10 insect species for at least 10 years. Additionally, studies must have either been restricted to a protected area or span a gradient of land use types (e.g., from developed to protected); and by “protected” we mean relatively isolated from land conversion rather than any legal or political designation.

Analyses of Northern California butterfly data involved visualization of population trends averaged at the site level, estimation of population trends at the species level, calculations of changes in mean elevation of occupancy per species, and ordination of inter-annual population variation in association with natural history traits. Full details on all methods are given in *SI Appendix* material, but in brief our visualization of populations (in Fig. 1C and D) was based on z-transformed probabilities of observation that we have shown to be indices of abundance (57). Estimation of coefficients summarizing population change over time (insets in Fig. 1C and D and shading of points in Fig. 2) is based on hierarchical Bayesian binomial models as presented in previous work with this data (41). Changes in average elevation per species (Fig. 1E-G) used sample-(or visit-) based rarefaction to impose an equal number of simulated visits to a site in repeated resampling to calculate differences between time windows. The specifics of time windows were motivated by a desire to understand change before, during and after a millennium drought (2011-2015) that was the single most impactful climate event (during our records) on the montane populations. Finally, redundancy analysis (RDA) combined many lines of information into one picture of population-specific change over time with respect to population-specific traits (Fig. 2).

## Acknowledgements

Thanks To David Wagner for organizing the Entomological Society of America symposium where much of this paper was originally presented. M.L.F was supported by a Trevor James McMinn professorship.

