## Supplemental material for "Insects and recent climate change"

**This file includes:**

Supplemental materials
Figure S1

### Supplementary Methods

***Literature survey of long-term studies.*** We surveyed the literature for long-term monitoring studies of insects using Web of Science and the following keywords: TS=(insect* OR lepidoptera* OR hymenoptera* OR diptera* OR hemiptera* OR coleoptera* ) AND TS=(climate OR weather) AND TS=("long term" OR "long-term" OR monitor*), which generated 2,246 citations from 1995 through 2019 (survey conducted in February of 2020). Shown in Table 1 is the subset of those studies that encompass at least 10 years of continuous sampling for at least 10 species of insects and include analyses of weather data potentially influencing insect populations.

***Population trends by site***. To facilitate visualization of population trajectories across species (Fig. 1C and D), we standardized observations among sites and among species, using z-score standardization on the probability of observation for a species within a site (specifically, the number of days observed divided by the number of visits to a site within a year), and then averaged z-scores across species to produce a mean probability of butterflies being observed (in units of standard deviations). The data being standardized (the ratio of positive observations to visits) has been shown previously to be a proxy for abundance (1). Visualizations (Fig. 1C and D) used data from all sites, and all species by site combinations (populations) that were present in at least five years.

***Calculation of changes in average elevation by species***. Elevational changes in population dynamics were examined by comparing the average elevation at which species were observed among the following three windows: 1977 – 1985, 2002 – 2010, and 2011 – 2019. The comparison between the first two windows goes from the earliest years that we have mountain data to right before the millennium drought years, and also mirrors a previously-published analysis (2). The comparison between the 2nd and 3rd windows encompasses before the drought years to the present, and then finally we compared the first and 3rd windows of time as our complete span of data. We used data from four sites for these analyses: Rancho Cordova (RC), Lang Crossing (LC), Donner Pass (DP), and Castle Peak (CP), thus reaching from the valley floor to the top of the mountain, but not including Washington (WA) because sampling there did not begin until 1988. A resampling (rarefaction) approach was used in the calculation of elevational shifts, to control for unequal sampling effort among sites over time. In brief, for each iteration of the rarefaction for a given time window, a sample (with replacement) of visits was drawn for each site (from the actual list of visits) to match the lowest sample size for any site in that year. A mean (rarefied) elevation was then calculated for each species across 1000 resampling iterations for a given time window, and a difference in average elevation was then calculated between time windows for each species. Results are shown both in Fig. 1E-G and also in Fig. S1 where space allows for the three histograms to be plotted on x axes with the same scale.

***Trait analyses of populations over time by elevation.*** Associations between species-specific traits and inter-annual variation in montane populations were examined using redundancy analysis (RDA). For each species at every site, the probabilities of observation, or fractional day positives (3), were calculated as the number of days with positive sightings divided by the number of visits per year and standardized as described above. Ordination was performed on the matrix of standardized detection probabilities, where each row in the matrix was a species of butterfly at a particular site and each column was the detection probability in a particular year. The predictor variables for the RDA were site residency (whether or not a species has maintained a local breeding population), overwintering stage, voltinism, geographic distribution, wingspan, a qualitative ranking of habitat generalism (from 1 to 5), sensitivity to annual average daily minimum temperatures, sensitivity to annual average daily maximums, sensitivity to annual precipitation, elevational range, a measure of upslope shift (from Fig. 1G), and weedy or ruderal status. The latter (weedy or ruderal status) describes more dispersive species often associated with exotic hosts (2). Sensitivity to climate was represented by beta coefficients from weather variables estimated with a hierarchical Bayesian binomial regression model run for each species, as described elsewhere (3). As in Halsch et al. (4), climate data used in the hierarchical models were based on 270m grid maps portraying average monthly values for minimum and maximum temperatures, and cumulative precipitation (5–7), extracted from cells that overlap monitoring sites, and averaged for water years (September through August). For visualization of temporal population patterns, points in ordination space (Fig. 2) were colored by a "trend" estimate, which was the coefficient associated with year from another round of hierarchical Bayesian models focused only on predicting day positives through time.

The redundancy analysis of montane population trends and species traits, performed on the covariance matrix using the vegan package (v.2.5-6) in R (8, 9), was focused on the west slope of the Sierra Nevada, specifically using data from four sites: Washington (WA), Lang Crossing (LC), Donner Pass (DP), and Castle Peak (CP) (Fig. 1B). Data were subset to the years

1988 to 2019 (1988 is the start of continuous sampling at Washington). Populations (site-specific occurrences of species) were only included in this analysis if they had been observed in at least ten years at a site. We are here treating populations (rather than species) as the unit of analysis, which builds on previous results that among-population heterogeneity of response to weather is large and comparable to among-species response (3).

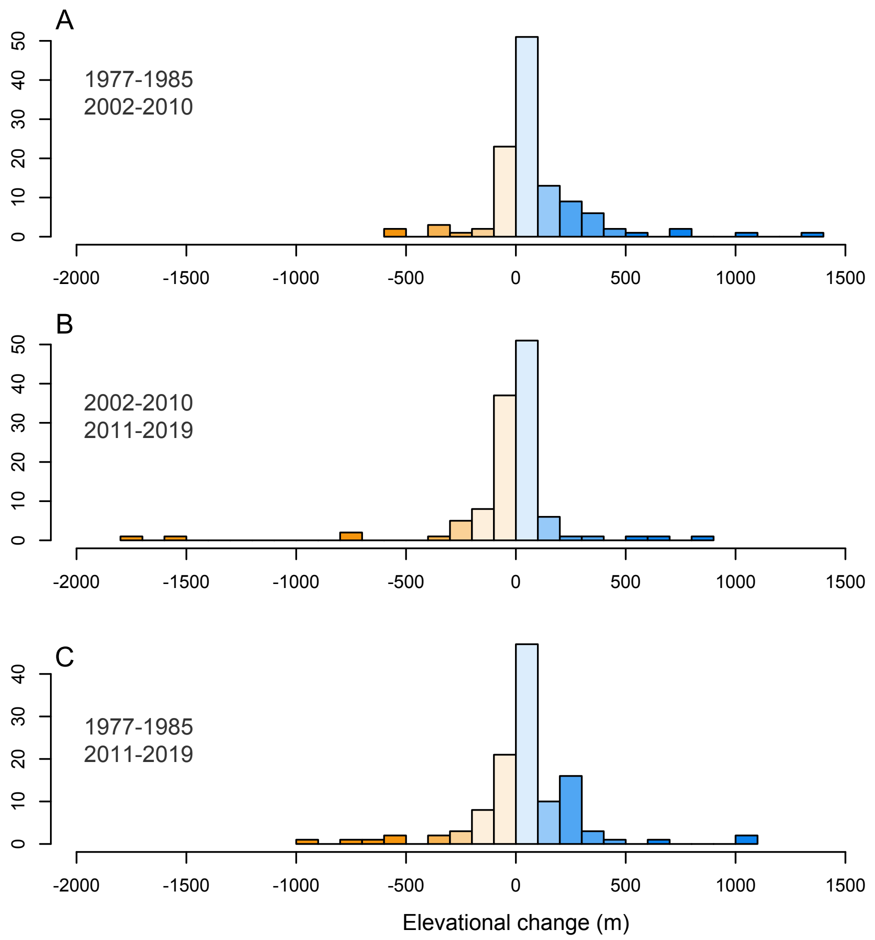


**Fig. S1.** Histograms of elevational shifts across species, or changes in the average elevation at which species were observed between different windows of time. The windows of time are as follows: 1977-1985 are the earliest nine years in the montane data; 2002-2010 are nine years immediately before the start of the millennium mega-drought in western North America; 2011-2019 are the most recent nine years of data (nine was used as the length of the windows so that the last two intervals would not be overlapping). For example, in the top panel, a species with a value of 500 is a case where the average elevation of observation for that species was 500 meters higher in 2002-2010 as compared to 1977-1985 (based on 1000 rarefied simulations to account for variation in sampling effort among sites). Colors in histograms are for visualization with darker orange corresponding to more negative (downward) shifts and darker blue being more positive (upslope) shifts. One-sample t-tests evaluate the hypothesis that each distribution has a mean different from zero, as follows: (*A*) $t = 3.82$, $p = 0.00022$, (*B*) $t = -1.59$, $p = 0.12$, and (*C*) $t = 1.63$, $p = 0.11$.
